# NF-kB c-Rel is dispensable for the development but is required for the cytotoxic function of NK cells

**DOI:** 10.1101/2021.01.19.427356

**Authors:** Y Vicioso, K Zhang, Parameswaran Ramakrishnan, Reshmi Parameswaran

## Abstract

Natural Killer (NK) cells are cytotoxic lymphocytes critical to the innate immune system. We found that germline deficiency of NF-kB c-Rel results in a marked decrease in cytotoxic function of NK cells, both in vitro and in vivo, with no significant differences in the stages of NK cell development. We found that c-Rel binds to the promoters of perforin and granzyme B, two key proteins required for NK cytotoxicity, and controls their transactivation. We generated a NK cell specific c-Rel conditional knockout to study NK cell intrinsic role of c-Rel and found that both global and conditional c-Rel deficiency leads to decreased perforin and granzyme B expression and thereby cytotoxic function. We also confirmed the role of c-Rel in perforin and granzyme B expression in human NK cells. c-Rel reconstitution rescued perforin and granzyme B expressions in c-Rel deficient NK cells and restored their cytotoxic function. Our results show a previously unknown role of c-Rel in transcriptional regulation of perforin and granzyme B expressions and control of NK cell cytotoxic function.

## Introduction

Natural Killer (NK) cells are innate immune cells with anti-tumor and anti-viral activity^1, 2^. These cells express a wide range of activating and inhibitory receptors that upon engagement by cognate ligands on target tumor cells regulate their anti-tumor activity^3, 4^. Upon activation, NK cells kill their target cells in a multistep-wise fashion which involves synapse formation between NK cells and target cells, expression of FAS ligand (Fas-L), and secretion of pro-inflammatory cytokines and cytotoxic granules that induce target cell death ^5, 6^. The granule exocytosis pathway is the primary mechanism used by NK cells to kill tumor and virus infected cells^7, 8^. Cytotoxic granule-mediated killing by NK cells largely depends on two proteins, perforin (prf1) and granzyme B (gzmb)^9, 10, 11, 12^. Perforin is a protein produced by both cytotoxic T cells and NK cells that form holes in the target cell membrane where granzymes enter to destroy the target cell^13, 14^. Granzymes are a family of serine proteases with gzmb being the most powerful pro-apoptotic member of this family^15, 16, 17^. Under pathological conditions such as in cancer, prf1 and gzmb are reported to be downregulated in NK cells compromising its cytotoxic function leading to NK cell exhaustion.^18, 19, 20, 21^. Thus, it is imperative that we identify key factors that regulate cytotoxic function in NK cells. Previously, we reported that inhibition of GSK3 signaling led to increased NK cell cytotoxicity via NF-κB activation^22^.

The Nuclear Factor-κB family of transcription factors consist of five subunits: p50, p52, RelB, p65, and c-Rel. Of these, c-Rel is predominantly expressed in the cells of myeloid and lymphoid lineages^23^. c-Rel global knockout (c-Rel^-/-^) mice exhibit comparable morphology and behavior to that of wild type (WT) mice, however c-Rel^-/-^ mice have profound immune system defects^24, 25, 26^. One such defect is in the thymic development of Foxp3^+^ regulatory T-cells^27, 28^. In addition, B cells from c-Rel^-/-^ mice exhibit proliferation and immune-specific functional defects.^29^ Specifically, c-Rel-deficiency impairs B cells ability to produce antibodies and T-cells ability to produce the cytokines, IL-2, IL-3, GM-CSF, TNF-α, and IL-6^26, 30^. It was recently shown, that B and T lymphocytes from a patient with homozygous mutation that eradicate c-Rel expression exhibit impaired T and B cell activation^31^. However, the role of c-Rel in NK cells remains poorly defined.

To address this, we used three different models including germline c-Rel knockout mice (c-Rel^-/-^), NK cell specific c-Rel conditional knockout mice (c-Rel^f/fcre^) and c-Rel shRNA transduced primary human NK cells. We show that c-Rel deficiency impairs NK cell anti-tumor activity in vitro and in vivo suggesting c-Rel as a key regulator of NK cell anti-tumor function. Finally, we show that c-Rel regulates prf1 and gzmb expression by directly binding to their promoters and c-Rel reconstitution reverses functional defect in c-Rel^-/-^ NK cells by restoring prf1 and gzmb expression. This study reveals a new role for c-Rel in NK cell-mediated anti-tumor immunity.

## Results

### NK cell maturation is unaltered in c-Rel^-/-^ mice

c-Rel deficiency impairs the generation and maintenance of activated regulatory T cells and reduces activated B and T cell proliferation and survival^24, 27, 29^. However, the role of c-Rel in NK cell has not been studied in detail. First, we analyzed c-Rel expression in NK cells compared to T and NKT cells. Flow cytometry analysis showed that murine NK cells express c-Rel comparable to T and NKT cells (Fig. 1a). NK cells develop from NK progenitor (NKp) cells, which originated from Common Lymphoid Progenitor (CLP) cells (Fig.1b). We confirmed c-Rel expression in NKp cells as well as differentiated CD27^+^ and CD11b^+^ NK cell subsets (Fig.1c). To determine whether c-Rel plays a role in NK cell development and maturation, we compared the total percentages and numbers of NK cells in germline c-Rel knockout mice (c-Rel^-/-^) compared to WT mice. Bone marrow (BM) and spleen (SPL) did not show any significant changes (Fig. 1d, e), however, reduced percentages of NK cells were observed in the lymph nodes (LN) (.34% to .23%) and the liver (4.7 to 2.6%) of c-Rel^-/-^ mice (Fig. 1d). c-Rel-deficiency also led to reduced total NK cell numbers in the blood (0.47×10^6^ to 0.2×10^6^), LN (0.03×10^5^ to 0.01×10^5^), and liver (0.09×10^5^ to 0.03×10^5^) (Fig. 1e). In mouse, four NK cell subsets are defined based on their maturation stages (Fig. 1b). Analysis of these four NK cells subsets in bone marrow and spleen of c-Rel^-/-^ mice did not show any major differences in percentages (Fig. 1f) or absolute numbers (Fig. 1g) compared to age/sex matched WT mice. In addition to CD11b, mature NK cells express markers such as DX5 and CD43. No measurable differences were found in the expression of these markers indicating that NK cell maturation process in c-Rel^-/-^ mice, is similar to that of WT mice (Fig. 1h).

**Figure 1.**
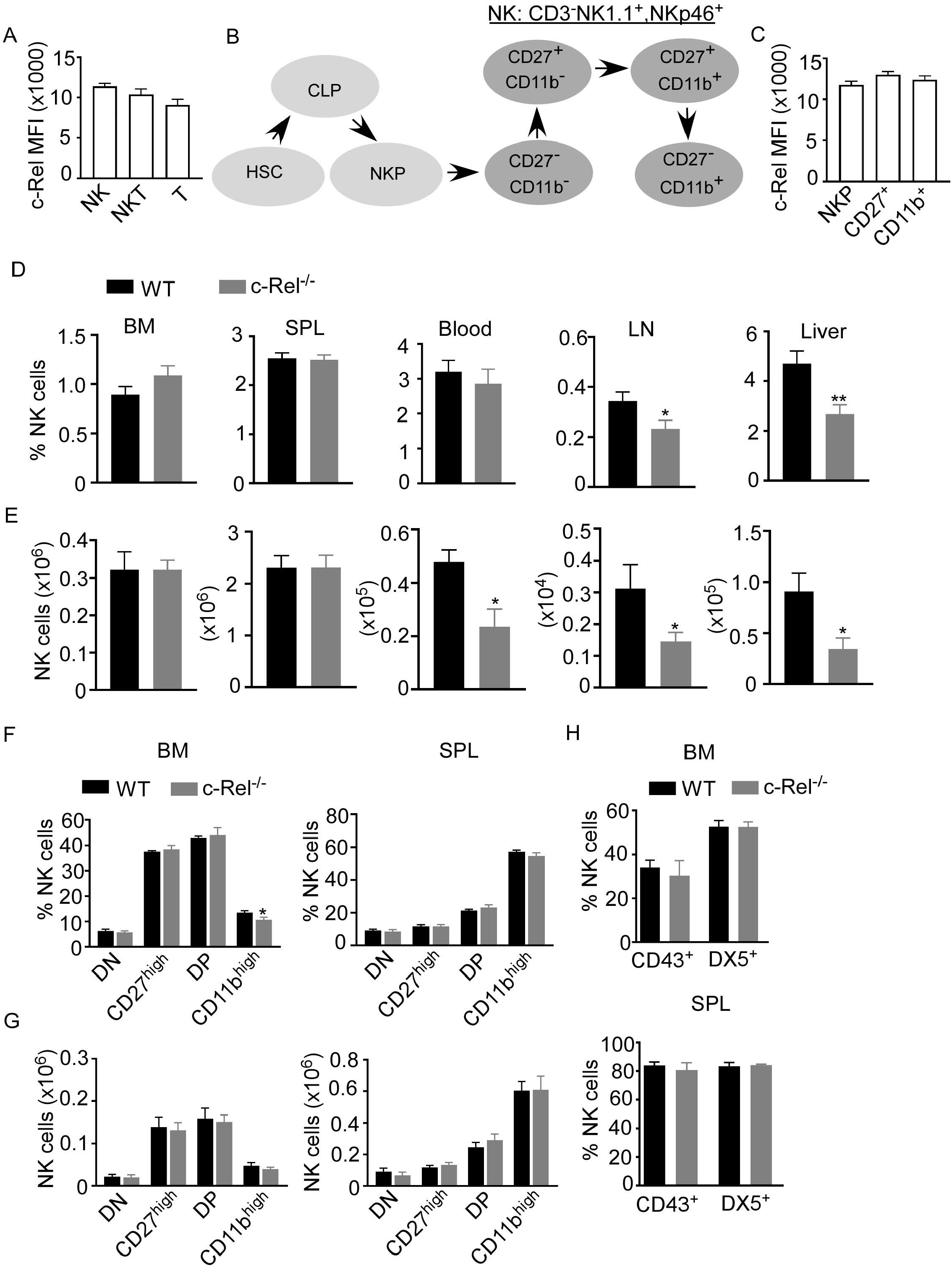
c-Rel is expressed in mouse NK cells. Total splenocytes from WT mice were stained intracellularly for c-Rel and analyzed via flow cytometry. a, Summary data of c-Rel expression on gated CD3^-^NK1.1^+^ (NK), CD3^+^NK1.1^+^ (NKT), and CD3^+^NK1.1^-^ (T) splenic cells. b, Schematic of NK cell development: hematopoietic stem cell (HSC) commit to common lymphoid progenitor (CLP) cells, which give rise to NK progenitors (NKp). NK cell maturation stages are immature (CD27^-^, CD11b^-^) and (CD27^+^, CD11b^-^), mature (CD27^+^, CD11b^+^), and terminally differentiated NK cells (CD27^-^, CD11b^+^). c, Summary data of c-Rel expression on gated CD3^-^ NK1.1^-^CD122^+^ NK progenitors (NKp), immature and transient CD3^-^NK1.1^+^CD27^+^ NK cells, and mature CD3^-^ NK1.1^+^CD11b^+^ NK cells. Each column (a,c) represents the mean ± SEM of data analyzed from 4 mice (n=4). Isotype IgG served as negative control for staining. d, Percentages of gated CD3^+^NK1.1^+^ NK cells in WT and c-Rel-deficient (c-Rel^-/-^) mice in the indicated organs. e, Total numbers of NK cells from WT and c-Rel^-/-^ mice. CD3^-^NK1.1^+^ NK cells were analyzed for the expression of CD27 and CD11b maturation markers. f, Percentages of double negative (DN), CD27^high^, double positive, and CD11b^high^ cells gated on CD3^-^NK1.1^+^ cells from the indicated organs of WT and c-Rel^-/-^. g, Total numbers of NK cell subsets from WT and c-Rel^-/-^ mice. Each column represents the mean ± SEM of 13 mice. For all flow cytometry analysis live cells were gated using forward (FSC-A) and side scatter (SSC-A). Total numbers of NK cells per organ were derived by using the formula [(NK% of total cells/100) x total cell number per organ)] and NK cell subsets numbers were determined using the following formula [(NK cell subset % of total NK cells/100) x total NK cell number per organ)]. For bone marrow analysis femurs and tibia of mice were collected (2 of each/mouse). h, Summary data of maturation markers CD43 and DX5 expression in gated NK1.1^+^CD3^-^ cells from the indicated organs (n=3). All data represents mean ± SEM. Data were analyzed by unpaired Student’s t-test. *p<0.05, ** p<0.01.

### Cytotoxic function is inhibited in NK cells from c-Rel^-/-^ mice

As we did not observe any significant changes in NK cell maturation in c-Rel^-/-^ mice, next we assessed if c-Rel is required for NK cell cytotoxic function. We performed calcein-AM release assay to test in vitro cytotoxicity of NK cells from WT and c-Rel ^-/-^ mice. Figure 2a shows that depending on the NK cell to target ratio, NK cells from WT mice killed 44-70% of murine 8093 acute lymphoblastic leukemia (ALL) cells compared to only 14-24% killed by c-Rel^-/-^ NK cells. Interestingly, this is not specific to 8093 tumor cells. WT NK cells killed 39-85%, while c-Rel^-/-^ NK cells killed only 17-40% of murine B16F10 melanoma cells (Fig. 2a). Cytokines such as IL-2 and IL-15 are involved in the activation of NK cells^32, 33^. Hence, we treated both WT or c-Rel^-/-^ NK cells with IL-2 or IL-15 for 24 hours and analyzed cytotoxicity. As shown in figure 2b, cytokine treatment did not enhance cytotoxicity in c-Rel^-/-^ NK cells. We then, tested if c-Rel-deficiency impairs NK cells killing *in vivo*. Melanoma tumor cells were implanted subcutaneously in WT recipient mice and tumor volume was monitored using calipers. Once tumors reached 50mm^3^, mice were randomized to be treated on day 5 and day 7 either with WT NK cells, c-Rel^-/-^ NK cells, or PBS at the indicated time points (Fig. 2c). Figure 2d shows that two injections of WT NK cells inhibited tumor growth, while c-Rel^-/-^ NK cells treated tumors failed to do so. This signifies a key role of c-Rel in NK cell cytotoxic function.

**Figure 2.**
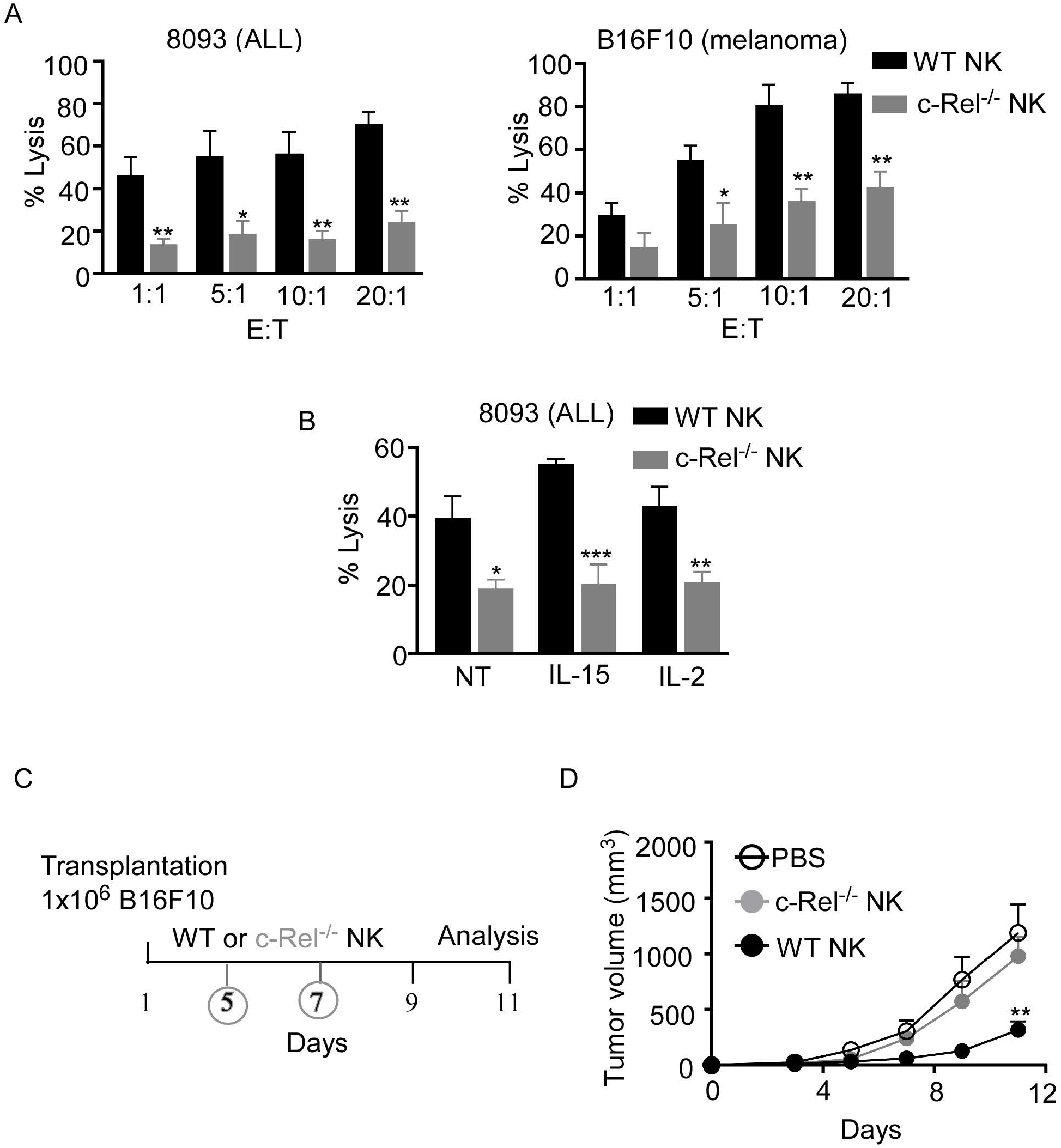
c-Rel deficiency impairs NK cell anti-tumor function. Cytotoxicity of NK cells from WT and c-Rel^-/-^ mice was evaluated using calcein AM release assay. a, Percent lysis of Acute Lymphoblastic Leukemia (ALL) and melanoma (B16F10) murine tumor cells co-cultured with either WT or c-Rel^-/-^ NK cells for four hours at the indicated effector (E) target (T) ratio. Tumor cells alone was used as negative control and tumor cells lysed with detergent served as positive control. b, Percent lysis of ALL cells after four-hour co-culture with either WT or c-Rel^-/-^ NK cells pre-treated with media (NT), IL-15 (100 ng/mL), IL-2 (1000 U/mL) for 24 hours. Data represents mean ± SEM of 2 or more independent experiments with NK cells pooled from the spleen and bone marrow of 2 mice of each genotype (n=4). c, Schematic of in vivo melanoma subcutaneous tumor model in WT background, treated with either PBS (negative control), WT NK cells or c-Rel^-/-^ NK cells on day 5 and day 7. d, Tumor volume of WT mice treated with either PBS, WT NK cells or c-Rel^-/-^ NK cells at the indicated time points. Data represents mean ± SEM of two independent experiments (n=5 per group). Data were analyzed using unpaired Student’s t-test. *p<0.05; **p<0.01; ***p<0.001.

### NK cell specific conditional c-Rel knockout mice exhibit NK cell dysfunction

Given that, c-Rel-deficiency has detrimental effects in B, T, and antigen presenting cells, it is crucial to dissect the cell intrinsic versus the extrinsic effects of c-Rel deficiency in NK cells. To understand how c-Rel-deficiency affect NK cell maturation and function in a cell intrinsic manner, we generated NK cell specific c-Rel-deficient mice (c-Rel^fl/fl^Ncr1^Cre^) by crossing c-Rel-flox to Ncr1^iCre^ mice. Western blot analysis shows deletion of c-Rel in purified NK cells from c-Rel^fl/fl^Ncr1^Cre^ mice compared to controls c-Rel^fl/fl^ mice (Fig. 3a). We then performed flow cytometry analysis of total NK cell population and found that compared to c-Rel^fl/fl^ control mice, c-Rel^fl/fl^Ncr1^Cre^ mice exhibit reduced percentages of NK cells in the spleen (2.8 to1.5%) and a slight reduction in the bone marrow (0.9 to 0.8%) (Fig. 3b, c). Analysis of CD11b and CD27 expression on NK cells, reveals increased percentages of CD27 single positive NK cells (40 to 55%), but a reduction of double positive (41 to 33%), and a reduction of CD11b^+^ (15 to 11%) NK cells in the bone marrow of c-Rel^fl/fl^Ncr1^Cre^ mice compared to c-Rel^fl/fl^ mice (Fig. 3d, e). Interestingly, NK cell subsets percentages were not altered in the spleen (Fig. 3e). Similar to c-Rel^-/-^ mice, c-Rel^fl/fl^Ncr1^Cre^ mice showed a reduction in the percentages of NK cell in the blood (3.3 to 1.8%), LN (0.38 to 0.26%), and liver (5.3 to 3.8%) compared to age/sex matched c-Rel^fl/fl^ mice (Fig. 3f). Lastly, we tested whether c-Rel deficiency impairs NK cell ability to kill tumor cells in vitro and found that indeed, c-Rel^fl/fl^Ncr1^Cre^ NK cells kill only17% of B16F10 tumor cells compared to 27% killing by NK cells from c-Rel^fl/fl^ control mice (Fig. 3g).

**Figure 3.**
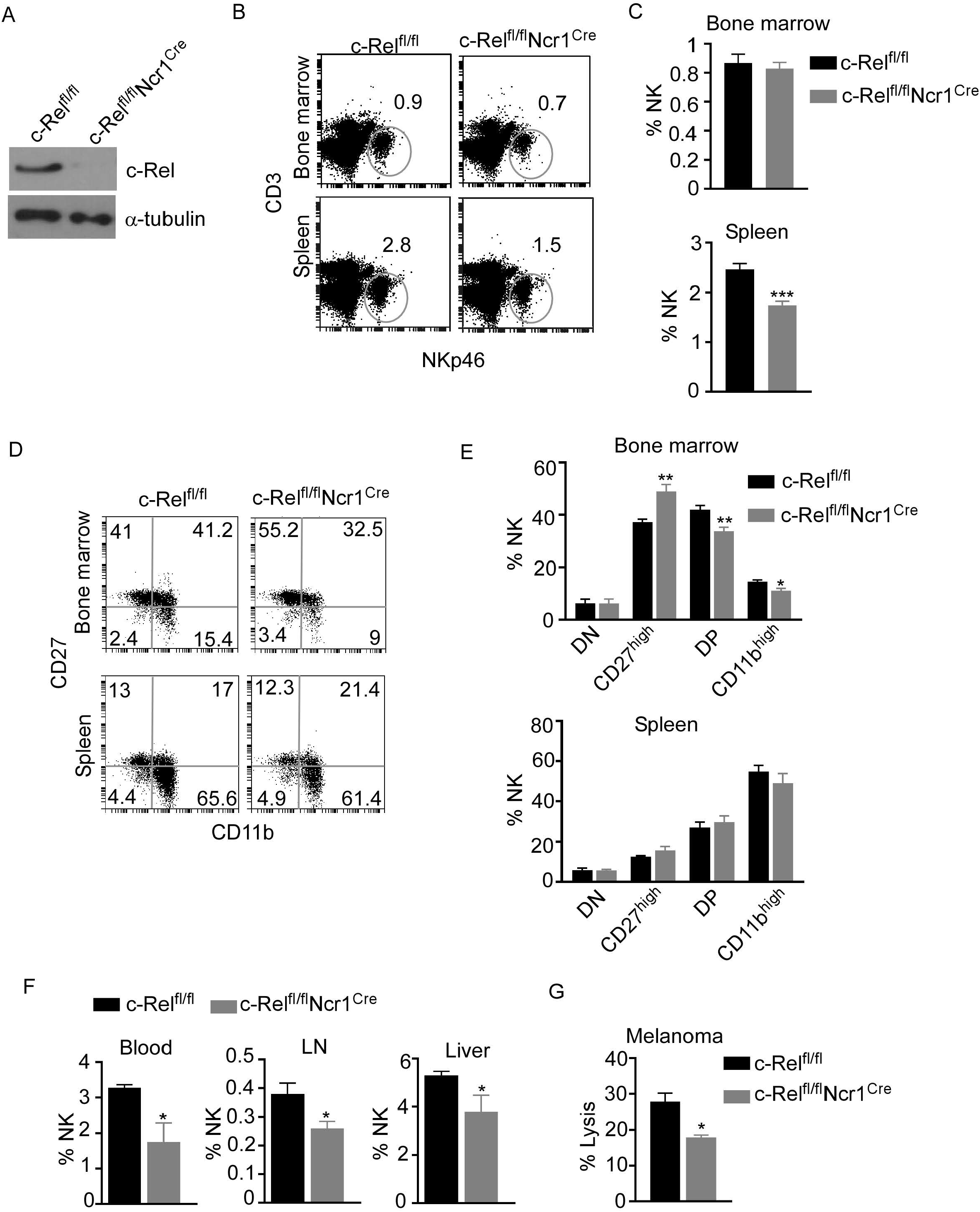
Loss of c-Rel alters cell-intrinsic NK cell maturation and anti-tumor function. NK cell specific c-Rel deficient (c-Rel^fl/fl^Ncr1^cre^) and c-Rel flox (c-Rel^fl/fl^) mice were used to evaluate the cell-intrinsic effects of c-Rel deficiency in NK cells. a, Western blot analysis of total lysate of uncultured NK cells pooled from the bone marrow and spleen of c-Rel^fl/fl^ and c-Rel^fl/fl^Ncr1^re^ mice (n=4 mice per group). Alpha tubulin was used as loading control. b, Representative plots and c, Summarized data of flow cytometry analysis CD3^+^NKp46^+^ NK cells from the bone marrow and spleen of the indicated mice. d, Representative plots and e, Summarized data of CD27 and CD11b expression in bone marrow and splenic CD3^-^NKp46^+^ NK cells. Each column represents the mean ± SEM of 5-8 mice (n=5-8). f, Percentages of NK cells in the indicated organs of c-Rel^fl/fl^ and c-Rel^fl/fl^Ncr1^cre^ mice (n=4). g, Percent lysis of melanoma tumor cells after co-culture with either c-Rel^fl/fl^ or c-Rel^fl/fl^Ncr1^cre^ NK cells for 4 hours as measured by LDH release assay. Data represents mean ± SEM of technical replicates. Data were analyzed using unpaired Student’s t-test. *p<0.05; **p<0.01; ***p<0.001.

### c-Rel deficiency impairs cytotoxic function in human NK cells

Up to this point, we have shown that c-Rel-deficiency impairs murine NK cell’s anti-tumor function in a cell-extrinsic as well as cell-intrinsic manner. However, murine NK cells differ phenotypically from human NK cells and findings cannot simply be extrapolated from one system to the other. To determine if c-Rel play a role in human NK cell function, we first, evaluated c-Rel expression in primary NK cells from healthy donors. In accordance with our analysis of c-Rel in murine NK cells, human NK cells also express c-Rel at a comparable level to that of NKT and T cells (Fig. 4a, b). We then used shRNA to ablate c-Rel expression in NK92 human NK cell line. Western blot analysis of NK92 cells transduced with shcRel shows around 50% reduction of c-Rel expression compared to shGFP control cells after 72 hours of viral tranduction (Fig. 4c). This reduction is also seen at the mRNA level (Fig. 4d). We then analyzed cytotoxicity of these cells using four different human tumor cells as target cells. As shown in figure 4e, c-Rel knockdown significantly impairs in vitro tumor killing by human NK cells. We also tested the effect of reducing c-Rel expression in primary human NK cells. To do this, we used the c-Rel inhibitor pentoxifylline^34, 35, 36^. Figure 4f shows that PTXF treatment in primary NK cells leads to a reduction of c-Rel protein expression, without affecting NF-κB p65 expression. Next, we tested in vivo cytotoxicity of NK cells pre-treated with or without PTXF using melanoma (WM164) xenograft tumor model (Fig.4g). Figure 4h, shows that four intratumor injections of control NK cells (pre-treated with water) significantly slows melanoma tumor growth from day 9-17 compared to PBS injected tumor bearing mice. NK cells pretreatment with PTXF significantly reduced this tumor inhibitory property of NK cells (Fig.4h).

**Figure 4.**
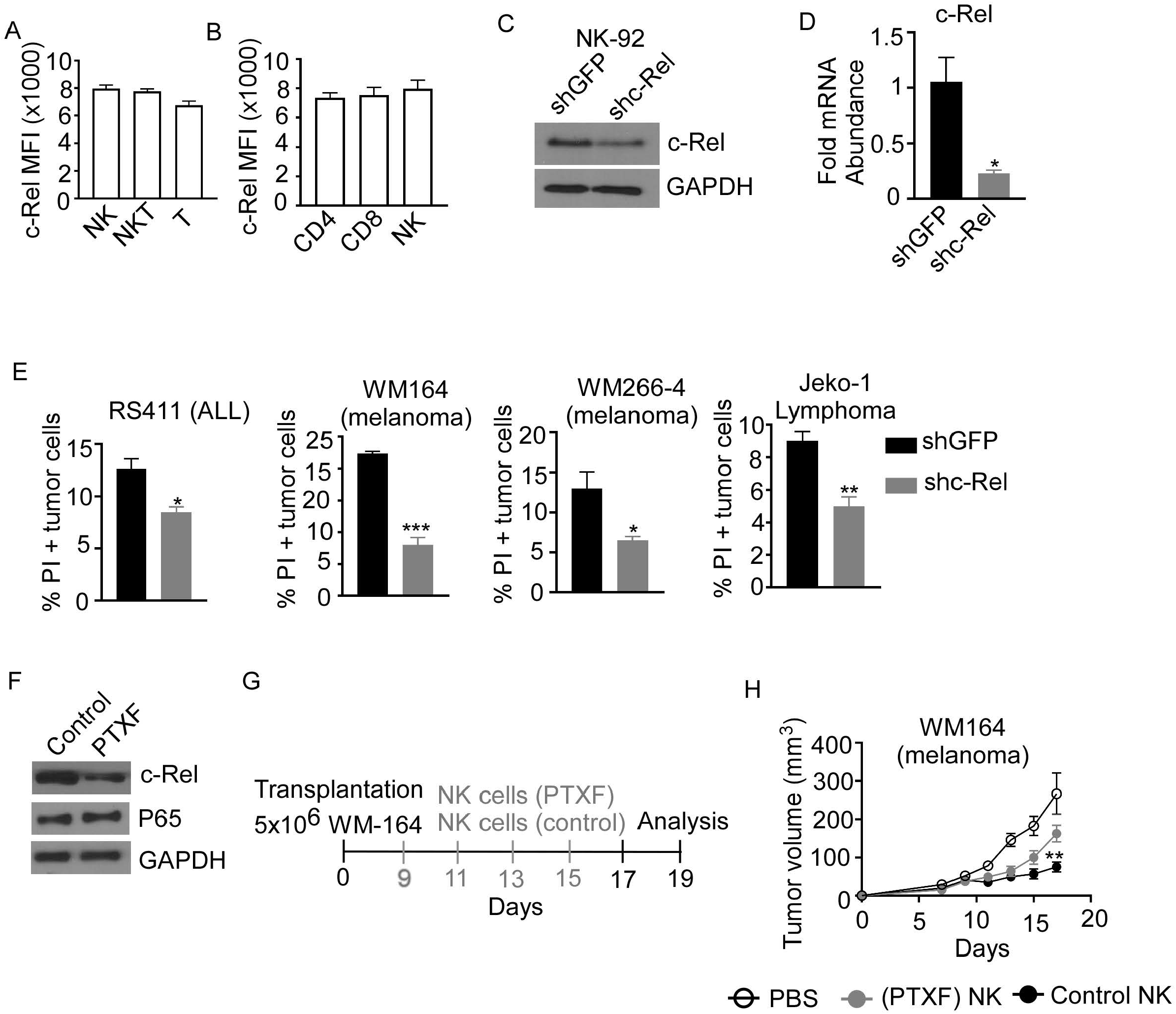
c-Rel regulates human NK cells anti-tumor function. c-Rel expression in primary human NK cells was evaluated by intracellular staining for c-Rel followed by flow cytometry analysis. a, Summary data of Mean Fluorescence Intensity (MFI) of c-Rel expression on gated CD3^-^CD56^+^ (NK), CD3^+^CD56^+^ (NKT) and CD3^+^CD56^-^ (T) human cells. b, c-Rel expression in (CD3^+^ CD4^+^) or (CD3^+^CD8^+^) T cells compared to NK cells. Each column represents the mean ± SEM of data collected from peripheral mononuclear cells (PBMCs) from 3 healthy donors. The human NK cell line (NK92) was transduced with shRNA lentiviral vectors containing either GFP (shGFP), a negative control or shRNA targeting c-Rel (shc-Rel) and analyzed for western blot or PCR. c, Western blot analysis on total cell lysates for c-Rel expression 48 hours post puromycin selection. GAPDH was used as loading control. Experiment done 3 times. d, mRNA fold change expression of c-Rel in NK92 cell 48 hours post puromycin selection relative to ribosomal RNA 18s. e, Percentage of Propidium iodide (PI) positive tumor cells after 4 hour co-culture with either shGFP or shc-Rel transduced NK92 cells, measured via flow cytometry. Data represents mean ± SEM of three technical replicates. f, Western blot analysis of c-Rel expression in primary NK cells treated with water (vehicle control) or Pentoxiphyline (PTXF) inhibitor (500 ug/mL) for 16 hours. GAPDH was used as loading control. Data is representative of 3 different experiments using NK cells from 3 healthy donors. g, Schematic of in vivo human melanoma (WM164) xenograft tumor model in NSG mice treated with either water pre-treated NK cells or PTXF pretreated NK cells for 16 hours prior to intratumor injections. h, Tumor volume of NSG mice treated with either PBS (negative control), water pretreated NK cells (positive control) or PTXF pre-treated NK cells (16 hours). Four total treatments were administered every other day starting on day 9 post tumor inoculation (10×10^6^ NK cells/injection). Data represents mean ± SEM of two independent experiments (n=5-6 per group). Data were analyzed using Student’s t-test. *p<0.05; **p<0.01; ***p<0.001

### c-Rel-deficiency renders NK cells defective in cytokine production

NK cells recognize tumor cells via their activating and inhibitory receptor repertoires on cell surface. We evaluated the expression of activating and inhibitory receptors in c-Rel^-/-^ NK cells compared to WT NK cells. Table 5a shows comparable expression of the activating receptors: NKG2D, Ly49D, DNAM1, Ly49C-H, CD244, CD16 and comparable expression of the inhibitory receptors: KLRG1, Ly49A, and LIR/PIR in c-Rel^-/-^ NK cells and WT NK cells at steady state. Following recognition and target engagement, NK cells kill target cells mainly via the prf1 and gzmb lysis pathway, but also through ligands that bind to death receptors in tumor cells such as fasL^6, 7, 8^. Flow cytometry analysis shows that c-Rel^-/-^ NK cells express fasL at a level that is comparable if not more than the WT NK cells (Table 5a). c-Rel is known to binds to the promoters of IFNγ and GM-CSF^23, 37^ We also measured cytokine production by WT NK cells compared to c-Rel^-/-^ NK cells upon activation. Figure 5b shows that activation of NK cells with IL-15, IL-2, or melanoma tumor cells induces the release of pro-inflammatory cytokines, TNF-α, GM-CSF, and IFNγ in WT NK cells. Compared to WT NK cells, c-Rel^-/-^ NK cells release less TNF-α, GM-CSF, and IFNγ upon activation with IL-15, IL-2, and B16F10 tumor cells (Fig. 5b). The immune suppressive cytokine IL-10 is slightly increased in c-Rel^-/-^ NK cells upon IL-15 and B16F10 tumor cell activation compared to WT NK cells (Fig. 5b). This data shows that c-Rel-deficiency renders NK cells defective in their ability to produce cytokines after cytokine or tumor induced activation.

**Figure 5.**
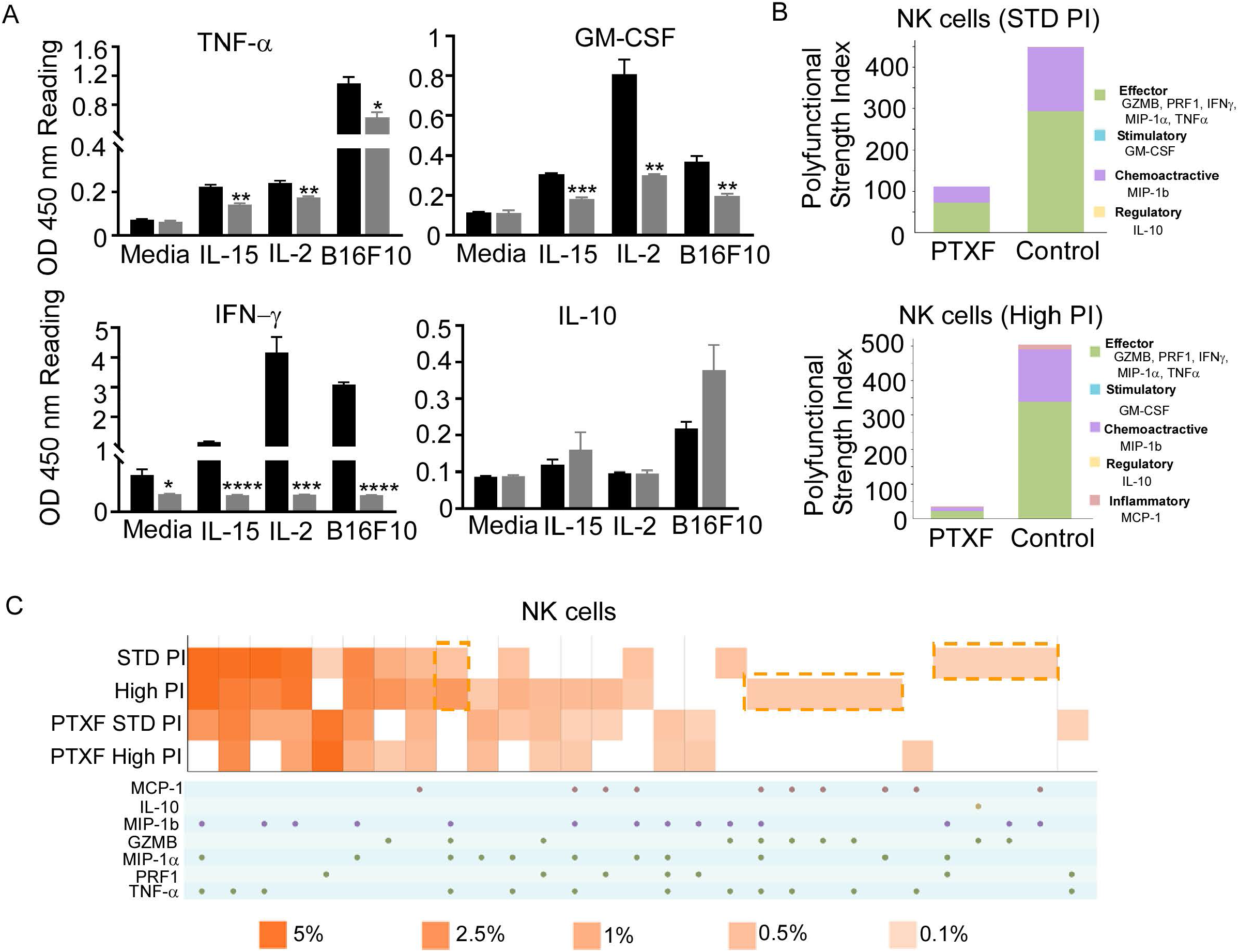
c-Rel deficiency impairs NK cell cytokine production. To evaluate the expression of receptors important for NK cell anti-tumor activity, single suspension cells were isolated from the bone marrow and spleen of c-Rel^-/-^ mice and analyzed by flow cytometry. a, Representative Mean Fluorescence Intensity (MFI) of gated NK cells expressing the indicated activating or inhibitory receptors from either the bone marrow or the spleen. Data represents one of 3 different experiments (n=3) with similar results. ELISA was used to evaluate the effect of c-Rel deficiency in NK cells ability to produce cytokines. b, Summary data from ELISA used to measure TNF-α, GM-CSF, IFNγ, and IL-10 in the conditioned media of WT or c-Rel^-/-^ NK cells cultured in media only (NT), IL-15 (100 ng/mL), IL-2 (1000 U/mL), or co-cultured with B16F10 melanoma murine tumor cells for 48 hours. NK cells were isolated from spleen and bone marrow of 3 WT and 3 c-Rel^-/-^ mice (n=3/genotype). Data represents mean ± SEM of technical triplicates. c, Polyfunctional Strength Index of the indicated cytokines/chemokines in healthy donor NK cells activated with IL-2 (10 ng/ mL), and either standard (STD,) or high PMA-neomycin (PI), following either water or PTXF (500 ug/ mL) treatment for 12 hours. d, Polyfunctional heat map illustrating single cell secretion of individual or combinations of cytokine/chemokines after the indicated treatments. Data were analyzed using unpaired Student’s t-test. *p<0.05; **p<0.01; ***p<0.001; ****p<0.0001.

To further confirm this, we performed a single cell cytokine analysis using control and PTXF treated human primary NK cells by Isoplexis. Polyfunctional Strength Index (PSI) is defined as the percentage of polyfunctional cells (secreting ≥2 cytokines) multiplied by the mean fluorescence intensity (MFI) of the secreted cytokines. Effector (GZMB, PRF1, IFNγ, MIP-1α, TNF-α), stimulatory (GM-CSF), chemoattractive (MIP-1b) and regulatory cytokines (IL-10) were measured at a single cell level. Here, the data show that activation with PMA/Ionomycin (PI) in control NK cells treated with water gives the expected high PSI, whereas treatment with the PTXF leads to lower PSI values (Fig. 5c). This inhibitory effect is greater with higher dose of PI stimulation, as shown in figure 5c. Figure 5d shows a heatmap visualization demonstrating that the positive PI stimulation of control NK cells gives the expected heterogenous heatmap visualization, with a higher frequency of polyfunctional cell subsets, while the PTXF treatment leads to a less heterogenous secretion profile, with a lower frequency of polyfunctional cell subsets. This effect was observed both in standard and high PI stimulation conditions.

### Reduced Granzyme B and perforin expression contributes to cytotoxic dysfunction in c-Rel-deficient NK cells

To determine the mechanism by which c-Rel regulates NK cell cytotoxicity, we evaluated the expression of prf1 and gzmb, two key cytotoxic mediators in NK cells^7, 9, 12, 14, 16, 38, 39^. PCR analysis of c-Rel^-/-^ NK cells shows reduced mRNA expression levels of gzmb (30%) and prf1 (50%) compared to WT NK cells (Fig. 6a). Since it has been shown that cytokines such as IL-2 and IL-15 can induce the expression of gzmb and prf1 in cytotoxic lymphocytes, we treated c-Rel^-/-^ and WT NK cells with either IL-2 or IL-15. Western blot analysis shows significant reduction of prf1 and gzmb protein levels in c-Rel^-/-^ NK cells treated with IL-2, or IL-15 (Fig. 6b). In this experiment we did not detected basal prf1 in NK cells, while gzmb showed reduced basal expression in c-Rel^-/-^ NK cells (Fig. 6b) compared to WT. We then asked whether this effect is unique to NK cells. PCR analysis of total pooled splenocytes and bone marrow cells shows a reduction in the mRNA levels of gzmb (51%) and prf1 (52%) in c-Rel^-/-^ cells (Fig. 6c). At the protein level, total cells show a reduction of both gzmb and prf1 in c-Rel^-/-^ cells compared to WT cells (Fig. 6d). PCR analysis in isolated NK cells from c-Rel^fl/fl^Ncr1^Cre^ and c-Rel^fl/fl^ control mice also showed a reduction in gzmb (40%) and prf1 (50%) mRNA levels indicating that c-Rel regulates gzmb and prf1 both intrinsically and extrinsically (Fig. 6e). Western blot analysis also shows reduced protein levels of gzmb and prf1 in NK cells from c-Rel^fl/fl^Ncr1^Cre^ mice compared to NK cells from c-Rel^fl/fl^ control mice (Fig. 6f). We then evaluated gzmb and prf1 expression in NK92 cell line after c-Rel shRNA mediated knockdown. Figure 6g and h, shows a reduction in the protein levels of gzmb (30%) and prf1 (40%) in NK92 cells transduced with shc-Rel compared to shGFP. mRNA levels of gzmb and prf1 were reduced by 70 to 80% (Fig. 6i). Finally, PTXF treated NK cells also showed reduced gzmb and prf1 expression correlating with reduced c-Rel in those cells, compared to control NK cells treated with water (Fig.6j). We then used Chip analysis to determine if c-Rel regulates gzmb and prf1 by directly binding to their promoters. Figure 6k shows enrichment at the gzmb and prf1 promoter in the c-Rel pull down compared to IgG control. This indicates that c-Rel binds the promoters of both gzmb and prf1. To determine whether c-Rel is sufficient to restore gzmb and prf1 expression in c-Rel^-/-^ cells, we transduced c-Rel^-/-^ pooled bone marrow and splenic cells (total cells) with either an empty or c-Rel containing lentivirus vector. Western blot analysis reveals that restoration of c-Rel expression in c-Rel^-/-^ total cells increases the expression of gzmb and prf1 (Fig. 6l). Finally, restoration of c-Rel restores c-Rel^-/-^ NK cell’s cytotoxicity against ALL murine tumor cells to the level of WT NK cells (Fig.6m).

**Figure 6.**
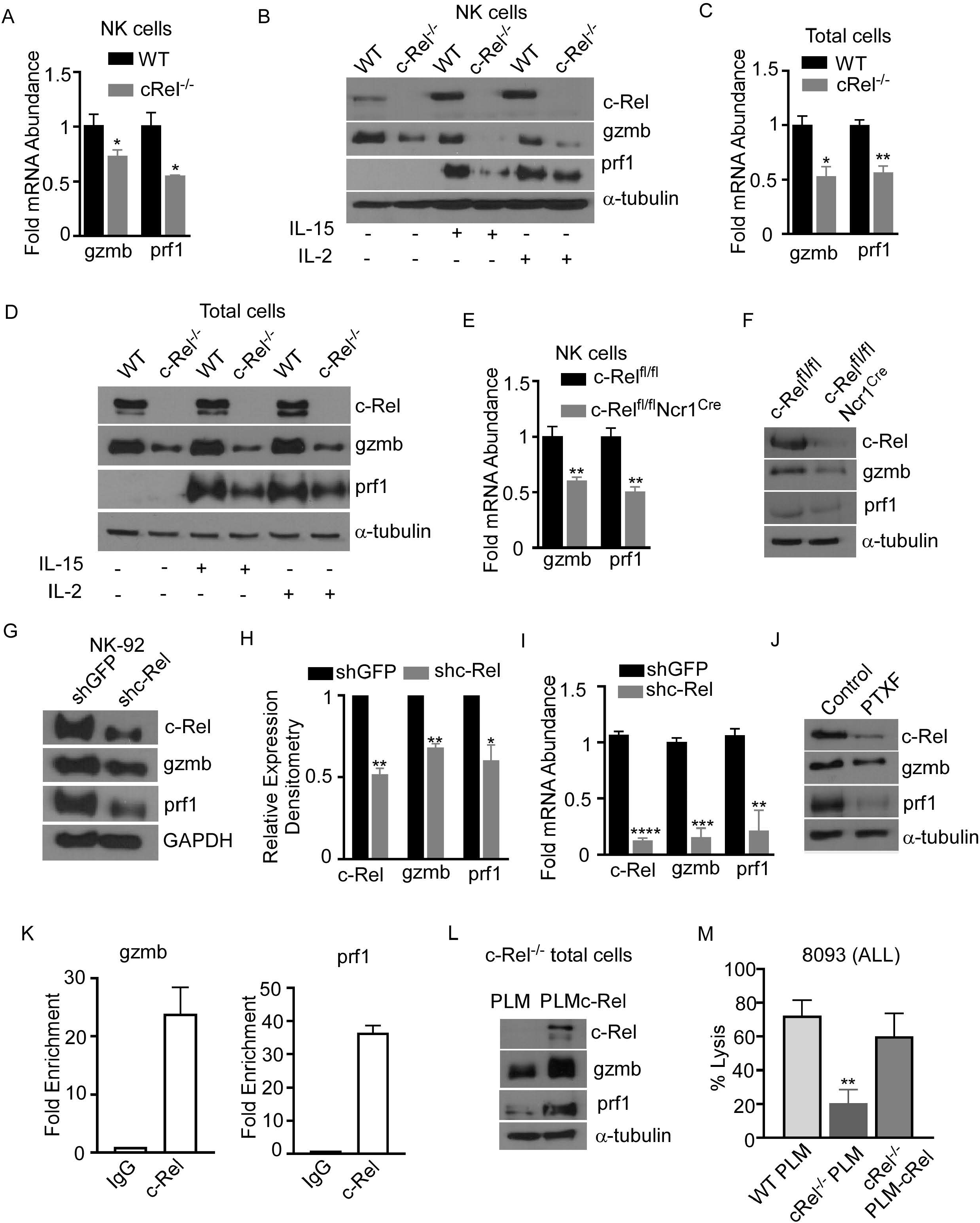
c-Rel regulates murine and human NK cell cytotoxicity via perforin and granzyme B. The expression of cytotoxic mediators granzyme B (gzmb) and perforin (prf1) was evaluated in NK cells pooled from the bone marrow and spleen of WT and c-Rel^-/-^ mice. a, PCR analysis of NK cells isolated from WT or c-Rel^-/-^ mice. Data represents fold change relative to GAPDH from at least 3 independent experiments n=3 mice per group. b, Western blot of WT and c-Rel^-/-^ NK cells treated as indicated for 48 hours. Experiment done with NK cells pooled from 3 mice per group (n=6). c, PCR analysis of gzmb and prf1 expression in total cells pooled from the spleen and bone marrow of WT or c-Rel^-/-^ mice (n=2 per group). d, Western blot analysis of total lysates from total cells pooled from the spleen and bone marrow of WT or c-Rel^-/-^ mice treated for 48 hours with the indicated cytokines (n=3-4). e, PCR of prf1 and gzmb in uncultured NK cells flow sorted from c-Rel^fl/fl^or c-Rel^fl/fl^Ncr1^Cre^ mice (n=2 mice per group). f, Western blot analysis of prf1 and gzmb in c-Rel^fl/fl^ or c-Rel^fl/fl^Ncr1^Cre^ NK cells enriched with MojoSort. The loading control used for western blot analysis was α-tubulin. Data represents at least 2 individual experiments. NK cells were purified from 2-3 mice per genotype. g, Western blot analysis and cumulative expression (h) of prf1 and gzmb in NK92 cells 72 hours after transduction with shGFP (empty vector) or shcRel (cRel targeting vector). NK92 cells were selected with 1ug puromycin for 48 hours prior to analysis. Data represents 2 independent experiments and GAPDH was used as loading control. i, PCR analysis of c-Rel, prf1, and gzmb expression in NK92 cells 72 hours post lentiviral transduction. Data represents mean ± SEM of three independent experiments with similar results. j, Western blot analysis of c-Rel, prf1 and gzmb in primary human NK cells 16 hours after treatment with PTXF. Water served as control. Experiment done 2-3 times with NK cells expanded from 2-3 different healthy donors. k, Chip assay of c-Rel binding to the prf1 and gzmb promoters in human primary NK cells. The DNA precipitated by c-Rel was detected by PCR. l, Western blot analysis of prf1 and gzmb of pooled bone marrow and splenic cells from c-Rel^-/-^ mice 72 hours after transduction with either empty lentiviral vector (PLM) or lentiviral vector containing WT c-Rel (PLM-c-Rel). Experiment done 2 times (n=2). m, Percent lysis of ALL cells by WT, or c-Rel^-/-^ NK cells 72 hours after transduction with PLM lentivirus compared to c-Rel^-/-^ NK cells transduced with PLM-WT-cRel lentivirus. Data represents 2 experiments done with NK cells isolated from 2-3 mice per genotype. Data represents mean ± SEM and were analyzed using Student’s t-test. *p<0.05; **p<0.01; ***p<0.001; ****p<0.0001.

## Discussion

In this study, we show that loss of c-Rel in mice leads to reduced numbers of NK cells in the peripheral organs: blood, lymph nodes, and liver. This data is in concordance with a recent report of immunologic profiling of blood from a six-year-old patient with complete loss of c-Rel expression showing increased numbers of CD4^+^, CD8^+^ T cells, but reduced numbers of CD19^+^ B cells and reduced numbers of CD3^-^CD56^+^ NK cells^31^. We did not observe any changes in total NK cell numbers or individual NK cell subsets in the bone marrow of c-Rel^-/-^ mice indicating that c-Rel is not essential for the development or maturation of NK cells within the bone marrow niche. Indeed, the expression of maturation markers DX5 and CD43 on NK cells isolated from c-Rel^-/-^ mice was also unaltered, compared to WT mice. This is not entirely unexpected as various studies in mice, report that c-Rel deficiency impairs T and B-cell proliferation and function without affecting the development and maturation of these cells^24, 25, 26^. Although c-Rel^-/-^ NK cells undergo normal maturation process, we provide evidence that loss of c-Rel impairs NK cell’s tumor killing function. We used ALL and melanoma cells as target cells representing liquid and solid tumors respectively. Tumor killing ability of c-Rel^-/-^ NK cells was significantly reduced against both target tumor cells regardless of the NK: Target ratio used. Given that IL-2 production is impaired in c-Rel^-/-^ mice, one could argue that the lack of IL-2 activation in c-Rel^-/-^ NK cells contributes to their cytotoxic dysfunction. However, this is unlikely as we show that exogenously added recombinant IL-2 or IL-15 fail to correct the NK cell dysfunction observed in c-Rel^-/-^ NK cells, suggesting that c-Rel regulates NK cell function in a cell intrinsic way. This is also evident in the B16F10 melanoma in vivo data as shown in figure 3d, where injecting WT NK cells into B16F10 melanoma tumors in a WT background led to tumor growth inhibition, while c-Rel^-/-^ NK cells had minor effect on tumor growth.

The stromal cell compartment of the bone marrow, as well as interaction with other immune cells have been shown to play a role in NK cell development and function^4, 40^. The functional defect of NK cells observed in c-Rel^-/-^ mice might be influenced by these extrinsic factors and to exclude this possibility, we developed NK cell specific c-Rel-deficient mice (c-Rel^f/fcre^). Though total percentage of NK cells in bone marrow remain unaltered, liver and lymph node showed reduced NK cells in both c-Rel^-/-^ and c-Rel^f/fcre^ mice. In consistent with c-Rel^-/-^ mice data, c-Rel^f/fcre^ mice also showed a reduction in NK cell cytotoxicity confirming cell intrinsic role of c-Rel in these cells.

Deficiency of c-Rel either by knockout or PTXF treatment resulted in reduced tumor cell killing by NK cells both in syngeneic and xenograft melanoma models respectively. PTXF inhibits anti-CD3 induced c-Rel expression and TNF-α production in T cells. A more recent study reports that c-Rel inhibition in T regulatory cells with PTXF enhanced melanoma tumor killing in a CD8 T cell-dependent manner^36^. Our data shows that PTXF treatment leads to reduced melanoma killing by NK cells. Reason behind these cell specific effect of c-Rel is not fully known. Recent reports show c-Rel playing an important role in regulatory T cells and myeloid cells checkpoint in cancer and inhibiting c-Rel as a method to enhance immunotherapy in cancer.^36, 41^ These observations suggest that the effects of PTXF or c-Rel inhibition differ among immune cell types and should be considered seriously while designing immunotherapies using c-Rel inhibitors.

NK cells recognize tumor cells through the direct interaction between its activating receptors with their ligands on tumor cells or through the lack of inhibitory receptor interaction with MHC-I on tumor cells.^42^ Of note, analysis of c-Rel^-/-^ NK cells revealed no alterations in the expression of activating and inhibitory receptors indicating that the capacity to recognize tumor cells is not affected by c-Rel deficiency.

Polyfunctional T cells are T cells producing multiple cytokines and are known to provide a more effective immune response than cells producing only a single cytokine.^43^ We analyzed and confirmed polyfunctionality of NK cells producing effector, stimulatory, chemoattractive and regulatory cytokines as indicated by polyfunctionality strength index (PSI). PSI was inhibited more than 4 fold in c-Rel inhibited human NK cells as shown in figure 4c, correlating to the reduced cytotoxicity observed in c-Rel deficient NK cells. NK cells kill tumor cells via release of cytotoxic mediators perforin and granzyme b or to a less degree via the interaction of NK cell FasL with the Fas receptor on tumor cells. We found comparable FasL expression in c-Rel-deficient NK cells compared to WT NK cells. However, a dramatic reduction in the mRNA and protein levels of perforin and granzyme B was noticed in c-Rel-deficient NK cells compared to WT. This was confirmed in both our c-Rel^f/fcre^ mice, and shRNA mediated c-Rel knockdown in human NK cells. It is noteworthy that expression of perforin and granzyme was not completely blocked in c-Rel^-/-^ NK cells, pointing to other transcription factors involved in the expression of these genes. As reported in previous studies, perforin basal level expression is minimal in NK cells, but can be activated with cytokines, IL-2, IL-15^39^. Interestingly, IL-2 or IL-15 treatment significantly increased perforin expression in WT NK cells, but not in c-Rel^-/-^ NK cells. Granzyme B levels were significantly reduced both at basal and cytokine induced conditions in the absence of c-Rel, pointing out to a differential transcriptional regulation of these genes by c-Rel. We show that c-Rel binds to both perforin and granzyme b promoters indicating a direct regulation of prf1 and gzmb by c-Rel. Rescuing c-Rel expression restores prf1 and gzmb expression and cytotoxicity in c-Rel^-/-^ NK cells, confirming NK cell dysfunction caused by c-Rel deficiency is mostly resulting from decreased perforin and granzyme B expression.

Our data show that c-Rel plays a cell intrinsic role in regulating human and murine NK cell anti-tumor cytotoxicity by regulating the expression of cytotoxic molecules prf1 and gzmb. It would be of interest to check c-Rel expression in NK cells from cancer or viral infected patients and analyze its contribution to the NK cell dysfunction observed in these patients.

## Methods

### Mice

Mice were bred at the Case Western Reserve University in accordance with the guidelines of the Institutional Animal Care Use Committee. The following strains (C57BL/6 genetic background) were used in this study: c-Rel^f/f^; (Purchased from The Jackson Laboratory), Ncr1^iCre^ purchased from Dr. Sexl^44^ (Institute of Pharmacology and Toxicology, Vienna, Austria), c-Rel germline knockout (c-Rel^-/-^) a generous gift from Dr. Ramakrishnan (Case Western Reserve University), and c-Rel^fl/fl^Ncr1^iCre/WT^ here called c-Rel^fl/fl^Ncr1^Cre^ were generated by backcrossing c-Rel^fl/fl^ to Ncr1cre at Case Western Reserve University. NOD SCID IL-2r Gamma (NSG) mice were purchased from Case Western Reserve University Athymic Animal Core. Experiments were conducted using age and gender-matched mice in accordance with approved institutional protocols.

### Cell isolation and stimulation

Bone marrow, spleen, blood, lymph nodes, and liver were harvested, and single cell suspensions were generated by pressing through sterile 40 uM nylon filter. RBCs were lysed and cells were washed in PBS containing 5% FBS (Sigma). Cells were stained for Flow cytometry analysis. For cell sorting experiments, bone marrow and splenocytes were surface stained with anti-NKp46, anti-CD3 and NKp46^+^CD3^-^ cells were sorted at the Case Western Reserve University Flow cytometry core using (BD Biosciences Aria) (purity>99%). For in vitro cytokine stimulations, MojoSort (BioLegends) purified NK cells (purity>80%) were plated at 1×10^6^ per mL and added either media alone, rmIL-15 (100 ng/mL), or rmIL-2 (1000 U/mL). Cells were harvested by pipetting with cold PBS after the indicated time and used for western blot and cytotoxicity assays.

### Flow cytometry and cell sorting

Cell surface staining of single-cell suspensions was performed using the following fluorophore-conjugated antibodies: c-Rel (sc-71) (purchased from Santa Cruz Biotechnology); CD3 (145-2C11), NK1.1 (PK136), IgG (MOCPC-21), IgG (HP6017), CD27 (3A10), CD11b (M1/70), CD43 (1B11), CD49b (DX5), NKG2D (CX5), Ly49D (4E5), DNAM-1 (10E5), Ly49 C-H (CF1H), NKG2A (16A11), CD244 (M2B4), CD16 (93), KLRG1 (2F1), Ly49A (A1), LIR/PIR (6C1), fasL (MFL3), NKp46 (29A1.4), CD3 (H1T3A), CD56 (51H11), CD4 (A161A1), CD8 (SK1) (purchased from BioLegend). Murine NK cells were defined as CD3^-^NK1.1^+^ cells. All data analysis was performed on viable cells by gating based on forward and side scatter. Flow cytometry was performed on Accuri 6C. Data were analyzed in Accuri 6C software analysis.

#### In vitro cytotoxicity assays

##### Calcein AM

NK cells were isolated from WT and c-Rel^-/-^ mice, or from c-Rel^fl/fl^ and c-Rel^f/f^Ncr1^Cre^ using MojoSort Magnetic Cell Separation (purchased from BioLegend). For anti-tumor cytotoxicity, 2-3×10^6^ NK cells were stimulated with the indicated cytokines or media alone for 24 hours prior to co-culture with tumor cells. The target cells were either the acute lymphoblastic leukemia cell line (8093) a kind gift from Nora Heisterkamp (City of Hope Comprehensive Cancer Center, Duarte, CA) or the melanoma cell line (B16F10) a kind gift from Dr. Bedogni (University of Miami, FL). Target cells were suspended in medium containing 5% FBS at a final concentration of 1×10^6^ cells/mL and incubated with 5uM calcein-AM (Life Technologies, Grand Island, NY, USA) for 1 hour at 37 °C with occasional shaking. After 3 washes with complete medium, 10,000 (B16F10) or 25,000 (8093) cells were plated in a 96 well plate. The number of NK cells was adjusted to achieve the different effector to target ratios. Cells were centrifuged and incubated at 37 °C for four hours, after which the plate was centrifuged and 70-100 μl of each supernatant was harvested and transferred into 96-well plates. Spontaneous (only target cells in serum free medium) and maximum release (target cells lysed using 0.1% Triton X-100) target cells were used as controls. Arbitrary Fluorescence Units (AFU) from each sample were measured using a Spectramax Gemini dual-scanning microplate spectrofluorimeter (Molecular Devices, Sunnyvale, CA, USA; excitation filter: 480±9 nm; band-pass filter: 530±10 nm). Percent lysis was calculated as described previously using the following formula: % specific lysis = 100 × (AFU mean experimental release-AFU mean spontaneous release)/ (AFU means maximal release-AFU means spontaneous release)^22, 45^. For the LDH-Glo cytotoxicity assay (Promega), 10,000 melanoma target cells were plated in a 96 well plate. After adding 10, 000 NK cells of each genotype, cells were centrifuged and incubated at 37 °C for four hours. After the incubation period the plate was centrifuged for 5 minutes at 1200 RPM and 5 μl of each supernatant was harvested and transferred into a 96-well plate. Spontaneous (only target cells in serum free medium) and maximum release (target cells lysed using 0.1% Triton X-100) target cells were used as controls. Samples were measured as specified by manufacture’s protocol using GloMax Discover Microplate Reader (Promega).

##### Flow cytometry-based cytotoxicity assay

In vitro killing of human tumor cell lines by NK92 cells was evaluated using flow cytometry as previously described^44, 45^. Briefly, tumor cells were resuspended in complete medium at a final concentration of 1×10^6^ cells/mL and labeled with 5 uM eFluor 670 (Thermo Fischer Scientific) as per protocol instructions. After washing, labeled target cells (1 × 10^5^) were added to 96-well plates along with indicated effector cells in complete RPMI media containing 100 U/mL of rhIL-2. Right before analysis 100ul of Propidium Iodide solution (Cell Signaling Technology) was added to the cells for 15 minutes at room temperature as indicated by manufacturer’s protocol. PI incorporation is used as a marker for late cell death as it intercalates with DNA in cells that have lost membrane integrity. Controls included target only wells (spontaneous death) and effector only wells to ensure that target cells only were in the eFluor 670^+^ gate. Results are shown as percent PI^+^ target cells ± SEM.

##### Cancer cell lines

The cancer cell lines mouse B16F10, human wm-164, and wm266-4 were a kind gift from Dr. B Bedogni (University of Miami, FL) and were cultured in DMEM supplemented with 10% FBS (Sigma) and 1% penicillin/streptomycin (HyClone). The murine acute lymphoblastic leukemia cell line (8093) was cultured in McCoy’s 5A media supplemented with: gentamycin (Gibco), 10% FBS (Sigma), Glutamax (Gibco), Sodium pyruvate (Gibco). Cells were washed every three days and supplemented with rmIL-3 (BioLegend), and BME (Gibco). Jeko-1 (purchased from ATCC) and RS411 cells (a kind gift from Dr. M. Dallas, Case Western Reserve University) were cultured in RPMI-1640 medium supplemented with 10% FBS (Sigma) and 1% penicillin/streptomycin (HyClone).

##### Tumor mouse models

Melanoma syngeneic tumors: B16F10 cells (1×10^6^ cells in 0.1 ml complete RPMI media) were injected subcutaneously (s.c.) into 8-week-old C57BL/6 mice, and mice were surveyed every other day. Tumors were allowed to reach 50-100mm^3^ and treated intratumorally with either purified WT, or c-Rel^-/-^ NK cells (2.5×10 /mouse) twice as indicated. For the melanoma (WM164) human in mouse xenograft: WM164 cells (5× 10^6^ cells in 0.1 ml complete RPMI media) were injected s.c. into 8-week-old NSG mice, and mice were surveyed every other day. We used pentoxifylline (PTXF) at 500 ug/mL to inhibit c-Rel in NK cells (Cayman Biochemicals). NK cells were pretreated with PTXF for 16-24 hours after which cells were washed to remove residual PTXF. PTXF or water control treated NK cells (10×10^6^) were injected in 100ul volume intratumorally at the indicated times. Tumor volumes were calculated as follows: v=1/2(Length × Width^2^)^46^ for the indicated times.

##### ELISAS

Supernatant were collected from the respective NK cells after treatment with either media only, IL-2, IL-15, or co-cultured with melanoma tumor cells (10: 1; NK: melanoma ratio) for 48 hours. NK cells were used at a final concentration of 1×10^6^/mL. Samples were stored at −80°C and thawed prior to use. For mouse TNF-α, IL-10, and GM-CSF ELISAs, 100 ul of undiluted sample per condition was used as specified by the manufacturer’s instructions. For the mouse IFNγ ELISA, 50 ul of undiluted sample per condition was used as indicated by manufacturer’s protocol. All the ELISA kits used were purchased from BioLegend. Binding of TNF-α, IL-10, GM-CSF and IFNγ was detected using secondary antibody, streptavidin-HRP, and TMB substrate solution (provided with specified ELISA kit). Substrate conversion was stopped after 20 minutes with 100 μL stop solution (2N H2SO4) purchased from BioLegend. Plates were washed with PBS plus 0.05% Tween20 in-between incubations. Assay diluent provided by the manufacturer or RPMI medium (Sigma) was used as negative controls, and specific standard proteins were used as positive controls. Standard reconstitutions and curves were generated as per manufacturer’s instructions for each assay. Optical density values were obtained using a microplate reader set to 450 nm (Bio-Rad iMark Microplate reader).

##### Single-cell barcode chip (SCBC) assay

To evaluate the production of cytokine/chemokines by NK cells following c-Rel inhibition, SCBC assay was performed in collaboration with IsoPlexis (IsoPlexis, Brandord, CT).^47^ Healthy donor PBMCs were treated with either water (negative control) or with PTXF (500 ug/mL) and stimulated with standard or high PMA-ionomycin (PI) and 10 ng/ mL human IL-2 overnight. Cells were then placed in a IsoPlexis’s IsoLight micro-chamber pre-coated with antibodies specific for cytokines and NK cell phenotypic markers. Secreted cytokines are capture and fluorescence is analyzed to measure the number of cytokines producing cells and the number of cytokines produced by single cell (polyfunctionality).

##### Human NK92 and primary NK cells culture

NK92 cells were a kind gift from Dr. Daniel Popkin (Innova Dermatology, TN, USA). Cells were cultured in alpha MEM media supplemented with: 2mM L-glutamine (Life Technologies), 1.5 g/L Sodium Bicarbonate (Fisher Chemical), 0.2 mM inositol (Acros Organics), 0.1 mM 2-mercaptoethanol (Life Technologies), 0.02 folic acid (Acros Organics), 12.5% horse serum (Sigma), 12.5% Fetal Bovine Serum (Sigma). Cells were washed and media was replaced every third day and supplemented with 200 U/mL IL-2 (PreproTech).

Human peripheral blood samples were collected from healthy donors in accordance with protocols approved by the Case Western Reserve University School of Medicine Human Studies Committee and Internal Review Board. Peripheral blood mononuclear cells (PBMCs) were obtained by using Ficoll-Paque and PBMCs were frozen in FBS containing 10% DMSO at a concentration of 10-20×10^6^ cells per vial. NK cells expansion was performed as previously described^45^, with minor modifications. Briefly,10 x10^6^ human PBMCs were cultured with 10 x10^6^ irradiated K562 clone 9 cell line (a kind gift from Dr. Dean A. Lee (Nationwide Children’s Hospital, Columbus, OH) for 2-3 weeks. Cells were grown in RPMI-1640 medium supplemented with 10% FBS (Sigma) and 1 % penicillin/streptomycin (HyClone). Fresh media was added every third day containing rhIL2 (100 U/mL; PeproTech). NK cells purity at the end of the 3-week coculture was (> 90%) as determined by flow cytometry (BD Accuri C6) using CD56 (5.11H11) and CD3 (HIT3a) antibodies from BioLegend.

##### Lentivirus production and infections

For shRNA mediated knockdown of c-Rel expression, NK92 cells were transduced with lentivirus vector pLKO-(Plasmid TRCN0000039984; Sigma), pLKO.1-puro eGFP shRNA control (Plasmid SHC005; Sigma) produced by packaging line HEK293FT. HEK293FT cells used for transfection were kindly provided by Dr. Ramakrishnan (Case Western Reserve University, Cleveland, OH). Cells were grown in DMEM supplemented with 10% fetal bovine serum (Sigma) at 37°C in a 5% CO2 incubator. HEK cells were transfected with pMD2.G (Plasmid 12259; Addgene); psPAX2 (Plasmid 12260; Addgene) in conjunction with pLKO.1 lentiviral vector targeting GFP or human c-Rel at a ratio of (1.5:3:4) using X-tremeGENE HP DNA transfection reagent (Sigma). 48h and 72h later viral particle containing media was harvested, clarified, and concentrated using Lenti-X concentrator (Takara) as per manufacturer’s protocol. Viral particle containing supernatant was added to NK92 cells growing in rhIL-2 containing media (200 U/mL) in 15 mL round bottom tubes and spun down at room temperature, 3480 RPM for 90 minutes. After spinfection virus particles were removed and replaced with complete alpha MEM media supplemented as above. 24 hours post transduction 1 ug/mL puromycin (InvivoGen) was added to the cultures and cells were harvested after 48 hours post-puromycin selection. NK cells were assessed for expression of c-Rel by either western blot or PCR and for cytotoxicity against different tumor cells types via flow cytometry.

For rescue experiments, the complementary c-Rel cDNA was cloned into the pLM-CMV-Hapuro-PL3 lentiviral plasmid^48^. Either bulk splenocytes plus bone marrow cells or isolated NK cells from WT or c-Rel^-/-^ mice were transduced via spinfection. Briefly, 100×10^5^ cells were plated in 96 well round bottom plates and 100 ul of viral particles was added. Cells and virus particles were spun at 22°C, 3480 RPM for 90 mins. After centrifugation, virus particles were removed, and cells were cultured in complete RPMI media.

##### Quantitative real-time PCR

NK cells from WT, c-Rel^-/-^, cRel^fl/fl^, c-Rel^fl/fl^Ncr1^Cre^ mice were either pre-enriched using (Mojosort, BioLegend) or flow sorted (BD Biosciences Aria) and RNA was isolated using RNeasy Plus Mini Kit (Quiagen) according to manufacturer’s protocol. For bulk bone marrow plus splenocytes or human NK92 NK cells, RNA was isolated using High Pure RNA Isolation Kit (Roche). From purified RNA, complementary DNA was synthesized using the Bio-Rad iScript Reverse Transcription Supermix cDNA synthesis kit (Bio-Rad). qrtPCR was performed in duplicate or triplicates in a 96 well plate using Bio-Rad iQ SYBR Green Supermix according to the manufacturer’s instructions. Data are represented as fold induction over the mean of control samples, using the ΔΔCt method. Primer sequences are as follows:

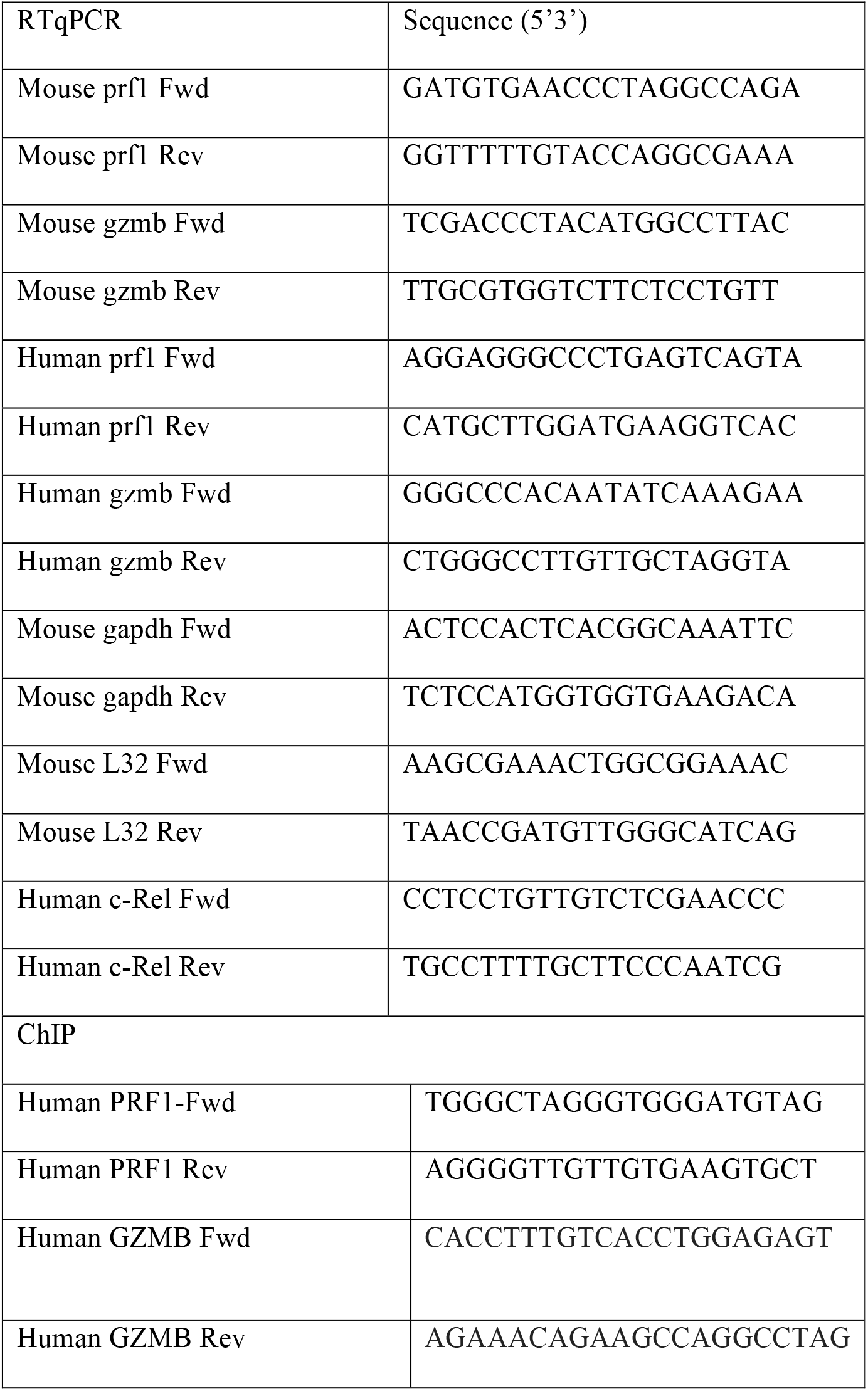

##### Immunoblots

For western blot analysis, 4 x10^6^ NK (Mojosort, purity >80%) or total combined splenocytes plus bone marrow from WT or c-Rel^-/-^ mice were stimulated for 48 hours in complete RPMI containing 10% fetal bovine serum (FBS) and 1% pen strep with either rmIL-2 (1000 U/ mL; BioLegend), rmIL-15 (100 ng/mL; BioLegend). Cells were washed with PBS and lysed with solubilization buffer containing 9 M urea. After sonication, lysates were spun down at 10,000 RPM for 5 minutes. Protein was quantified using (nanodrop nae, Case Comprehensive Cancer Center) and denatured with 4x lamely buffer followed by boil for 10 mins. 15-30 ug of total protein were loaded in 10% acrylamide gel. Proteins were transferred to nitrocellulose membranes (BioRad). Membranes were blocked with 5% dry milk in TBS containing 0.01% Tween and immunoblotted for mouse c-Rel (CD6), gzmb (sc-8022), p65 (sc-8008), GAPDH (sc-47724), prf1(sc-136994) purchased from Santa Cruz Biotechnology, mouse prf1 (3693s), human c-Rel (4727s) purchased from Cell Signaling, and alpha tubulin (cp06) purchased from Calbiochem. Mouse and Rabbit horseradish peroxidase (HRP) conjugated antibodies (Sigma) were used. Western blots were detected using chemiluminescence detection kit (Pierce) or chemiluminescence kit (BioRad).

### Chromatin immunoprecipitation (ChIP)

Analysis of c-Rel binding was performed as follows: 20×10^6^ expanded human NK cells were stimulated with IL-2 overnight. Cells were harvested, and proteins were crosslinked to DNA in two steps: (1) by adding DSG (Disuccinimidyl glutarate; ProteoChem) to a final concentration of 2mM for 45 minutes at room temperature followed by washing once with PBS and (2) by adding formaldehyde to a final concentration of 1.0% for 15 min at room temperature followed by the addition of 0.25 M glycine. Cells were washed twice with cold phosphate-buffered saline and resuspended in Farnham Lysis Buffer with the addition of protease inhibitor cocktail tablets (Sigma) followed by 15 minutes’ incubation with rotation at 4°C. Nuclear crude pellet was obtained by centrifugation at 4°C, 10,000 RPM for 10 minutes. Pellet was resuspended in RIPA (EMD Millipore) buffer and sonicated to shear chromatin by using the Sonicator 3000 Ultrasonic liquid processor optimized to 80 cycles of 8-s pulses with an output of 5 that gave a range of DNA fragments from 100 to 1,000 bp, as determined by 2% agarose gel electrophoresis.^49^ The soluble chromatin fraction was incubated with either anti-c-Rel antibody or anti-IgG control antibody (Biolegend) coupled to Protein G beads (Cell Signaling) overnight with rotation at 4°C. Immunocomplexes were washed with LiCl wash buffer and eluted with elution buffer. Protein-DNA cross-links were reversed at 65°C overnight with shaking. Samples were recovered by phenol/CHCl3/isoamyl alcohol extraction twice and purified as per kit instruction using Monarch PCR and DNA cleanup Kit (New England BioLabs). DNA was suspended in 200 ul pre-warm (55°C) elution buffer and stored at −80°C for real-time PCR amplification. The transcription factor binding site databases: Promo-ALGGEN and LASAGNA-Search 2.0 were used to find NF-κB or c-Rel specific binding sites in the promoter regions (10,000bp upstream and 1000 bp downstream) of perforin and granzyme B. Primers spanning the predicted NF-κB and c-Rel binding sites were designed using the Primer designing tool-NCBI-NIH. The following oligonucleotide sequences were used as primers to PCR amplify the region spanning the predicted site (atgggaaacta) NF-κB binding site in the granzyme b promoter: sense primer, 5’ CACCTTTGTCACCTGGAGAGT-3’ and antisense primer, 5’ AGAAACAGAAGCCAGGCCTAC-3’. For perforin, 5’TGGGCTAGGGTGGGATGTAG-3’ and antisense primer, 5’AGGGGTTGTTGTGAAGTGCT −3’ were used to amplify the region spanning the predicted site (agggaatggcca). iQSYBR green PCR analysis was performed using CFx96 Real-Time System. Identical thermocycler conditions were used as per manufacturer’s instructions. Each PCR was performed in triplicates. Since PCR amplification was being monitored by SYBRgreen, the absence of nonspecific amplification was determined by analyzing the dissociation curve of the PCR amplification products. Data collected were analyzed using Microsoft Excel and plotted on PRISM software. The amount of precipitated target sequence was calculated as the fold increase in signal relative to the background signal (Fold enrichment method, Thermo Fisher Scientific).

### Statistics

Data are shown as mean ± s.e.m. Two-tailed unpaired Student’s t test were used to determine significant differences (p< 0.05) using Prism software (Graph Pad, San Diego, CA). Sample size was not predetermined, but we used a number of mice that was consistent with prior experience with similar experiments.

## Data availability

Data supporting the findings of this study are available from the corresponding authors upon request. Materials will be provided with material transfer agreement in place as appropriate.

## Acknowledgements

This work was supported by NIH Award F31CA232648 (Y.V.), St.Baldrick’s scholar award (R.P.), Be positive (B+) Foundation grant (R.P.), 1R01AI116730-01A1 (P.R.). This research was supported by the Athymic Animal and Hemapoietic Biorepository and Cellular therapy core facilities, Shared Resources of the Case Comprehensive Cancer Center (P30CA043703) for their support. We thank the flow cytometry core and the radiation core facilities at Case Western Reserve University. We thank the department of Pathology for providing the Accuri C6 and other valuable resources to conduct this study. We also thank Dr. A. Tomaka for providing assistance with in vitro expansion of NK92 cells, Dr. Nora Heisterkamp for providing 8093 mouse ALL cells and Dr. T. de Jesus for providing technical assistance with chip assays. Finally, we thank Drs. A. Huang and J. Letterio, for insightful discussions.

## Contributions

Y.V performed the experiments, analyzed data and wrote the manuscript, K.Z. performed cloning and viral transductions, P.R contributed to study design, discussions, provided key reagents including c-Rel knockout mice and protocols, helped in troubleshooting and writing the manuscript. R.P. conceived the study, oversaw the study, designed experiments, guided research personnels, performed data analysis and wrote the manuscript.

